# An optimised protocol for quality control of gene therapy vectors using Nanopore direct RNA sequencing

**DOI:** 10.1101/2023.12.03.569756

**Authors:** Kathleen Zeglinski, Christian Montellese, Matthew E. Ritchie, Monther Alhamdoosh, Cédric Vonarburg, Rory Bowden, Monika Jordi, Quentin Gouil, Florian Aeschimann, Arthur Hsu

## Abstract

Despite recent advances made towards improving the efficacy of lentiviral gene therapies, a sizeable proportion of produced vector contains an incomplete and thus potentially nonfunctional RNA genome. This can undermine gene delivery by the lentivirus as well as increase manufacturing costs and must be improved to facilitate the widespread clinical implementation of lentiviral gene therapies. Here, we compare three long-read sequencing technologies for their ability to detect issues in vector design and determine Nanopore direct RNA sequencing to be the most powerful. We show how this approach identifies and quantifies incomplete RNA caused by cryptic splicing and polyadenylation sites, including a potential cryptic polyadenylation site in the widely used Woodchuck Hepatitis Virus Posttranscriptional Regulatory Element (WPRE). Using artificial polyadenylation of the lentiviral RNA, we also identify multiple hairpin-associated truncations in the analysed lentiviral vectors, which account for most of the detected RNA fragments. Finally, we show that these insights can be used for optimization of lentiviral vector design. In summary, Nanopore direct RNA sequencing is a powerful tool for the quality control and optimisation of lentiviral vectors, which may help to improve lentivirus manufacturing and thus the development of higher quality lentiviral gene therapies.

## Introduction

Recent years have seen considerable advances in the field of lentiviral gene therapy, with several clinical trials delivering stable, long-term transgene expression to treat disease (1)(2). However, several challenges remain before lentiviral gene therapies can be routinely implemented in the clinic. These include that a substantial proportion of lentiviral RNA can be non-functional. As a consequence, the physical titre (number of virus particles) is typically much higher than the infectious titre (number of infectious virus particles) (3). Although a low functional titre is not unusual for lentiviruses (it has been estimated that as few as 1% of HIV virus particles are capable of infection (4)), this must be improved in order to facilitate cost-effective, large-scale manufacturing of lentiviral gene therapies. An important cause of nonfunctional lentivirus is the presence of incomplete lentiviral RNA, which can occur for a range of reasons such as anomalous splicing, premature transcription termination at cryptic polyA sites or insufficient processivity of the polymerase due to the length of the construct (3)(5). In addition to lowering the infectious titre, this may reduce therapeutic efficacy due to an incomplete transgene and can also pose a safety risk due to loss of safety elements such as chromatin insulators or other regulatory components designed to limit the potential for oncogene activation (6). These issues can be solved by mutating or removing problematic sites or by reducing the length of the vector itself (which will remove potentially problematic sites and increase transcriptional efficiency) (3)(5). It is therefore important to have an assay in place that can identify not only the abundance, but also the causes of lentiviral RNA truncation. RNA sequencing using Illumina (3) and Nanopore direct cDNA (5) approaches have both been used to demonstrate improvements to lentiviral vectors, and sequencing has also been applied to other therapeutics, such as mRNA vaccines (7). However, there is currently no clear consensus on which sequencing technology is most effective for quality control of lentiviral vectors, and current cDNA sequencing approaches which rely on reverse transcription (RT) of the lentiviral RNA can introduce bias (8). In contrast, Nanopore direct RNA sequencing is a relatively new technology that enables the direct sequencing of long, potentially full-length, native RNA. Thus, it may present an unbiased, faster and more efficient way to assess lentiviral vectors. Additionally, as sequencing starts from the 3’ end of the RNA, the ends of sequencing reads correspond directly to the 3’ ends of the physical RNA molecules which allows for precise identification of truncated vector sequences. It has also proven successful during the recent pandemic, with the high throughput sequencing of SARSCoV2 viral genome (9)(10)(11). Finally, other long-read approaches such as Nanopore and PacBio also have the potential to capture the entire lentiviral RNA in a single read, enabling analysis at a single-molecule level. To facilitate quality control (QC) of lentiviral vectors early in development, there is a need for a quick, efficient, and cost-effective sequencing assay. Thus, to determine the most appropriate sequencing technology for lentiviral vector QC, this study aimed to compare Nanopore direct RNA sequencing with cDNA sequencing approaches (from Nanopore and PacBio). We found that Nanopore direct RNA sequencing enables the identification of cryptic splice and polyA sites that contribute to RNA truncations. Modifications to the sequencing protocol were also considered to improve the proportion of vector reads and to identify incomplete RNAs that are not naturally polyadenylated. This enabled us to identify RNA hairpins as major contributors to truncation in the analysed lentiviral vectors and to estimate the percentage of full-length RNA. Finally, we show that this analysis allows us to implement potential optimisations of the lentiviral vector sequence for increasing vector integrity.

## Results

### Nanopore direct RNA sequencing is an ideal long-read RNA sequencing approach for lentiviral vector QC

In order to compare different approaches for QC of lentiviral vector integrity, we decided to perform an initial experiment with “Globin LV”, a gamma globin expressing vector that had shown low functional titres. In this vector, gamma globin is expressed from a β-globin promoter (jointly labelled “Transgene” in Figure 1) adjacent to regulatory elements (HS2, HS3 and HS4). In addition, a short hairpin RNA (shRNA) targeting human Hypoxanthine Guanine Phosphoribosyltransferase (HPRT) is expressed from a human 7SK RNA Pol III promoter (“7SK”). A 400 bp extended core element of the chicken hypersensitivity site 4 (cHS4) β-globin chromatin insulator (“insulator” in Figure 1) is inserted in the 3’LTR as a safety and anti-silencing element. This insulator was previously shown to contain a potent cryptic splice acceptor site, which was identified as the cause for clonal dominance in progenitor and myeloid lineages of patients in clinical trials (12)(13). RNA was isolated from concentrated and purified lentiviruses that were produced using stable producer cell lines, which was then subjected to sequencing by Nanopore direct RNA, Nanopore direct cDNA and PacBio cDNA sequencing approaches to compare the three technologies. Read numbers varied by technology, ranging from 491,270 reads for Nanopore direct cDNA to 273,029 reads for Nanopore direct RNA (both Nanopore methods were performed using minION flow cells) and 50,159 reads for PacBio cDNA sequencing. In all cases the percentage of vector aligned reads was relatively low (less than 30%; Supplementary Table 1). The remaining reads aligned to the human genome, indicating that most of the sequenced RNA originates from the producer cells rather than the lentivirus and has been co-purified during lentivirus purification and RNA isolation.

**Fig. 1.**
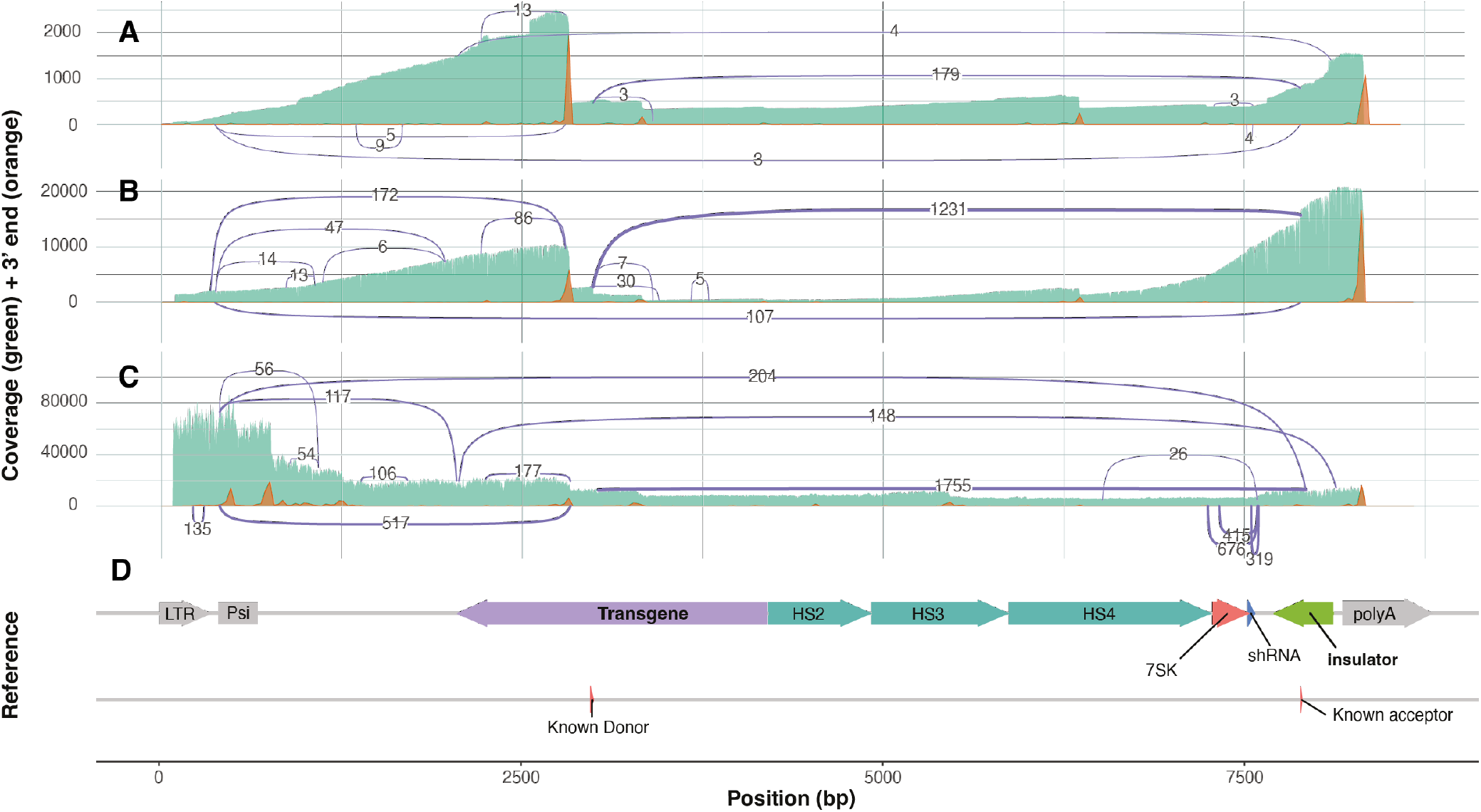
Comparison of three different long-read RNA sequencing approaches for lentiviral vector QC. Sequencing coverage (green), location of the 3’ ends of reads (orange) and splicing patterns (purple lines connecting splice donors and acceptors, with numbers indicating how many reads support each splicing event) for three different long-read sequencing technologies. **(A)** PacBio cDNA sequencing data. **(B)** Nanopore direct RNA sequencing data. **(C)** Nanopore direct cDNA sequencing data. **(D)** Reference sequence of the Globin LV, with various features annotated.

Sequencing coverage showed a step-like pattern of sharp changes which slowly decay from high to low coverage moving towards the 5’ end of the vector sequence in the PacBio cDNA and Nanopore direct RNA data (1, green track). This decay can be explained by reverse transcription (for PacBio cDNA) and sequencing (for Nanopore direct RNA) beginning at the 3’ end of the RNA, and not extending all the way to the 5’ end. Sharp changes in coverage, such as the one at 3000 bp, indicate sites leading to incomplete transcripts. Interestingly, the Nanopore direct cDNA data (Figure 1C) shows a different pattern: its coverage is highest at the 5’ end, with several sharp coverage changes that do not appear in the analysis of the other two approaches. These coverage changes correlate with internal polyA tracts in the vector reference sequence, suggesting that the 5’ coverage bias is due to internal priming during the reverse transcription step leading to the formation of fragments not reflecting the state of the lentiviral RNA. Plotting of the reads’ 3’ ends revealed the intended polyA site at the 3’ end of the reference sequence as well as three commonly used cryptic polyA sites (1, orange track) which coincide with many of the sharp coverage changes noted above. The most significant of these, located at 3000 bp was used by 36.82% of the reads in the PacBio cDNA sample (as compared to 33.62% of reads that terminate at the 3’ end of the vector) (1A) and 16.44% of reads in the Nanopore direct RNA sample (while 59.85% terminate at the 3’ end of the vector) (1B). Identification of cryptic polyA sites in the Nanopore direct cDNA data is complicated by the large number of internal priming sites, which also produces 3’ end peaks that are indistinguishable from those that originate from truncated RNA (Figure 1C). Importantly, numerous splicing patterns were identified in the Globin LV (Figure 1C, purple lines). While the Nanopore sequencing approaches picked up more splice events than the PacBio approach, likely due to the significantly higher number of reads enabling detection of rarer splicing patterns, several events were consistently observed across all three approaches. In particular, a splicing event between a previously identified splice donor site (data not shown) and the known splice acceptor site in the insulator (12)(13) had the highest number of supporting reads with all three sequencing technologies. This splicing event leads to the formation of an incomplete lentiviral RNA that contains the packaging signal (Psi), which allows these incomplete lentiviral transcripts to be packaged (14). These transcripts, however, lack a substantial portion of the therapeutic globin cassette and are therefore unlikely to confer a therapeutic benefit. Altogether, between the compared approaches, we found the Nanopore direct RNA sequencing technology to be the most appropriate for lentiviral RNA QC given the simplicity of the workflow and the comprehensive overview of identified splice events and cryptic polyA sites. While Nanopore direct cDNA sequencing provides more reads and higher coverage at the 5’ end of the vector, internal priming bias can severely confound identification of cryptic polyA sites. Similarly, for PacBio sequencing, the lower number of reads and high cost per base limited the identification of rarer splicing events.

### Nanopore direct RNA sequencing identifies cryptic polyA and splice sites

To further evaluate the utility of Nanopore direct RNA sequencing for lentiviral vector QC, we applied the technology for the analysis of four Wiskott-Aldrich Syndrome (WAS) lentiviral vectors, differing from each other only in their insulator sequences (described below). All four vectors express human WAS (“Transgene” in Figure 2) from a strong synthetic promoter. The 3’UTR consists of a Woodchuck Hepatitis Virus Posttranscriptional Regulatory Element (WPRE) that is commonly included in lentiviral vectors to increase expression (15). As for the Globin LV, a second cassette expresses an shRNA to target HPRT under the control of a 7SK promoter. All WAS vectors contain a 650 bp version of the cHS4 insulator (16), with a core sequence identical to that of the 400 bp cHS4 insulator containing the same known splice acceptor site (12)(13). While the first vector “WAS LV1” harbors an unmodified 650 bp insulator (in reverse orientation), the insulator sequence was modified in the three other vectors aiming to inhibit splice acceptor activity. Vectors “WAS LV2” and “WAS LV3” contain two and three A-to-T point mutations in “AG” splice acceptor motifs, respectively. Mutated in both vectors are the known splice acceptor site (12)(13) also present in the Globin LV, and an additional splice acceptor site predicted in silico (NetGene2 (17)), located outside the core region and thus specific to the 650 bp insulator version. Vector “WAS LV3” contains an additional mutation in a third “AG” motif located only 10 bp away from the known problematic splice acceptor site (12)(13), consistent with a published vector containing the same two point mutations in the cHS4 insulator core (13). A fourth vector “WAS LV4” contains an inverted cHS4 650 bp insulator (i.e. in forward orientation) with the purpose of de-coupling splicing and polyadenylation, processes which can co-stimulate each other (18). In this vector, the known and predicted splice acceptors are in the opposite orientation relative to the lentiviral polyA signal. Between 445,076 and 743,182 reads were generated for each sample, of which approximately 1.5-3% aligned to the vector reference sequence (Supplementary Table 2). This is much lower than for the Globin LVs and can be explained by the differences in purification process used. RNase treatment to degrade unpackaged, co-purified RNAs did not lead to an improvement in retrieval of lentiviral RNA (Supplementary Figure 1; Supplementary Table 3).

**Fig. 2.**
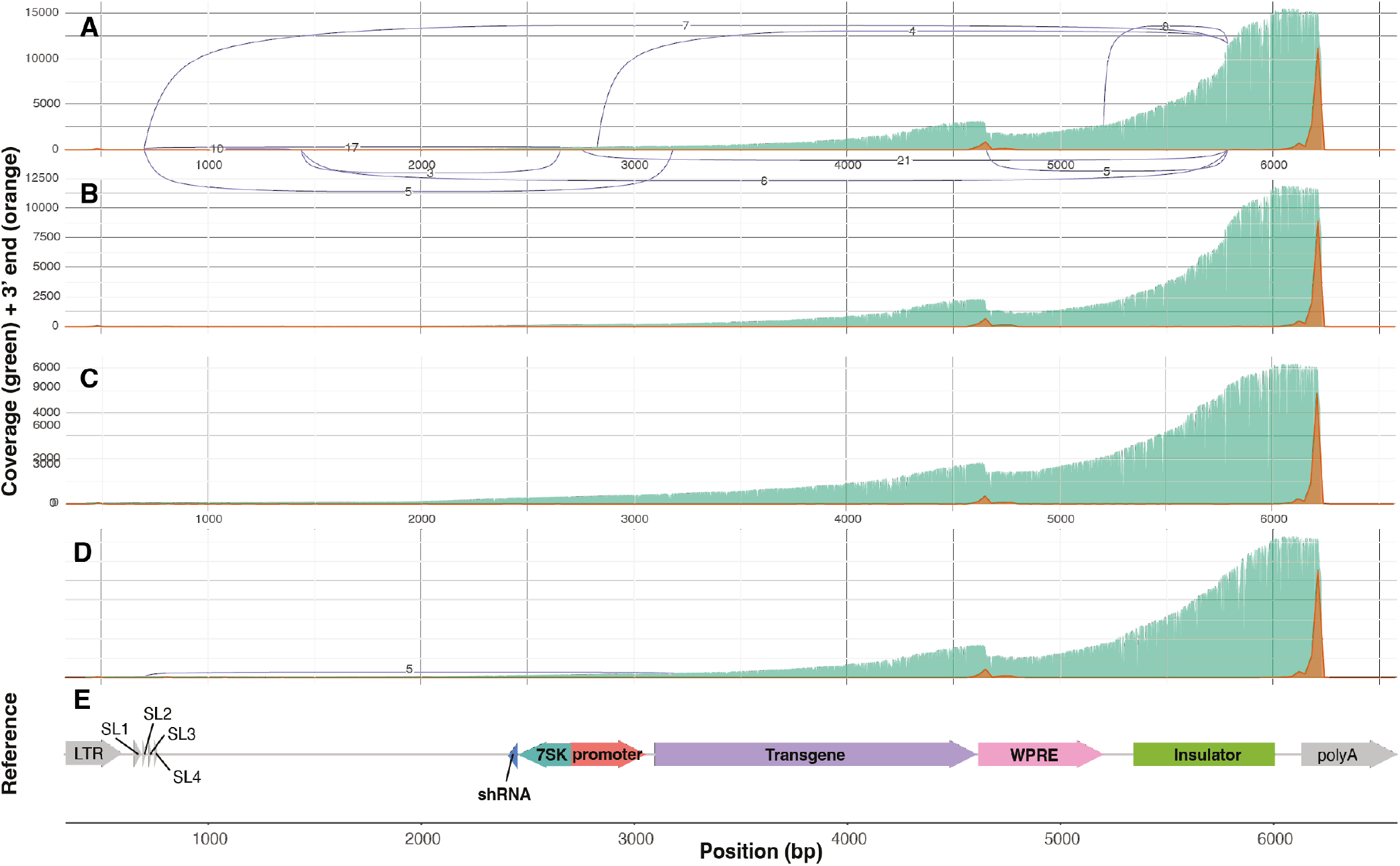
Using Nanopore direct RNA sequencing to assess Wiskott-Aldrich Syndrome vectors. Plots showing the sequencing coverage (green), location of the 3’ ends of reads (orange) and splicing patterns (purple lines connecting splice donors and acceptors) for four different Wiskott-Aldrich Syndrome vectors sequenced by Nanopore direct RNA sequencing: **(A)** WAS LV1, **(B)** WAS LV2, **(C)** WAS LV3, **(D)** WAS LV4. **(E)** Reference sequence of the Wiskott-Aldrich Syndrome vector, with various features annotated.

The sequencing coverage of WAS vectors (green, Figure 2) shows a strong pattern of 3’ to 5’ decay, with fewer sharp changes than for the Globin LVs, and a higher percentage of full-length vector (Figure 1). Analysis of the location of 3’ read ends of WAS vectors showed that while 80% terminated in the expected position at the end of the reference sequence (tall orange peak, Figure 2), 10% terminated around 4600 bp, in close proximity to the end of the WAS coding sequence (small orange peak, Figure 2). This coincides with a sharp change in coverage that occurs at the same position consistently across all four vectors, and we hypothesised that this might correlate with the presence of a cryptic polyA site. In addition to the polyA site at the 3’ end of the vector, the canonical polyA signal motif, AATAAA, was identified at four additional locations within the vector, although none were near 4600 bp. Instead, the only polyA motif identified near this site was the non-canonical ATTACA motif, approximately 10-20 bp upstream of where the reads terminate, which could potentially lead to premature polyadenylation. Alternatively, given that the putative polyA site is located close to the end of the WAS coding sequence and the large proportion of RNAs other than the lentiviral RNA in the sample (purple arrow, Figure 2E), we investigated whether these transcripts may represent endogenous WAS transcripts expressed from the cells used for lentivirus production. Endogenous WAS differs from the vector sequence by its 265 bp 3’ UTR, as well as by two single nucleotide variants due to two point mutations introduced in the coding sequence of the transgene. None of the reads terminating at 4600 bp aligned to the WAS 3’ UTR and the two point mutations were observed in 99% and 97% of reads across all samples, respectively. This indicates that the transcripts terminating at 4600 bp are being transcribed from the vector and terminate prematurely, potentially due to usage of the alternative 3’ ATTACA polyA site. Little splicing was observed in the WAS vectors, suggesting that splicing events are not a major cause of incomplete lentiviral RNA for these vectors (Figure 2). The original vector, WAS LV1 showed a small amount of splicing, with less than 100 out of 18,000 vector-aligned reads originating from splicing events. Most of these splicing patterns involved the known canonical acceptor site (12)(13), located in the chromatin insulator region (Figure 2), which was also observed in the Globin LV (Figure 1). As described above, we attempted to modulate splicing through modification of the vector either by introduction of point mutations in the splice acceptor or by inversion of the insulator. These modifications appear to be successful in reducing splicing into this site, as WAS LV2 and WAS LV3 showed no splicing at all, while WAS LV4 had only a single splicing event, involving a different acceptor site. This demonstrates the potential for Nanopore direct RNA sequencing to identify problematic sites in lentiviral vectors, to inform the design of optimized vectors, and to verify that introduced modifications reduce truncated lentiviral RNA.

### Artificial polyadenylation reveals additional sites of truncation

We then explored the use of artificial polyadenylation to capture incomplete lentiviral RNA that may not be naturally polyadenylated, such as those originating from incomplete transcription or those that are degraded or fragmented during library preparation. For this analysis, we chose the WAS vector with the wild-type insulator sequence containing the detected splice acceptor sites (WAS LV1). A second vector, WAS LV2 was also analysed using this method (see Supplementary Figure 2). Compared to the original, non-polyadenylated samples (Supplementary Table 2), the median read lengths of polyadenylated reads were decreased from 700 bp to 300 bp and a smaller percentage of them aligned to the vector (Supplementary Table 4). This can be explained by the abundance of co-purified short RNA species (e.g., rRNA, snRNA) that can be sequenced once polyadenylated.

Despite having fewer mapped reads, the polyadenylated samples show a similar pattern of sequencing coverage towards the 3’ end of the vector, whereas at the 5’ end coverage is increased (green, Figure 3), coinciding with the presence of sharp coverage changes and 3’ end peaks (orange, Figure 3B) in this region. These peaks, making up 11.26% and 8.61% of reads, respectively, are more pronounced than the one representing the potential cryptic polyA site close to the end of the WAS transgene (4.44% of reads), suggesting that these sites may play a significant role in the generation of incomplete WAS vector RNA. Upon closer examination of the vector reference sequence in these locations, both 3’ peaks occur around RNA stem-loop structures: the shRNA sequence targeting HPRT at around 2300 bp and several conserved stem-loops (SL1-SL4) present within the packaging signal (Psi) at around 500 bp (Figure 3C). It is possible that these RNA sites near stem-loops are being cleaved by endonucleases such as Drosha, which is known to recognise such structures(20). As demonstrated by these findings, artificial polyadenylation of lentiviral RNA before Nanopore direct RNA sequencing can facilitate the identification of a broader range of truncations. In addition to identifying additional types of incomplete lentiviral RNA, artificial polyadenylation enables a more precise estimate of the percentage of complete and incomplete RNA as all RNA fragments originating from the lentiviral vector should be captured. From counting the number of reads that terminate around the suspected cryptic polyA site (4.44%) and aforementioned stem-loop structures (19.88%), combined with the 1.31% of reads found to be spliced, we can estimate that at least a quarter of lentiviral RNA reads are incomplete in some way. In addition, approximately 15% of reads terminate prematurely, but not at cryptic polyA sites or known stem-loop locations (not in the 3’ end peaks in Figure 3). These may result either from insufficient polymerase processivity or instead correspond to lentiviral RNA that has been degraded or fragmented during the isolation and sequencing library preparation processes. Thus, we can estimate that approximately 60-75% of WAS LV1 RNAs are full-length, with 0-15% resulting from insufficient processivity of the RNA polymerase.

**Fig. 3.**
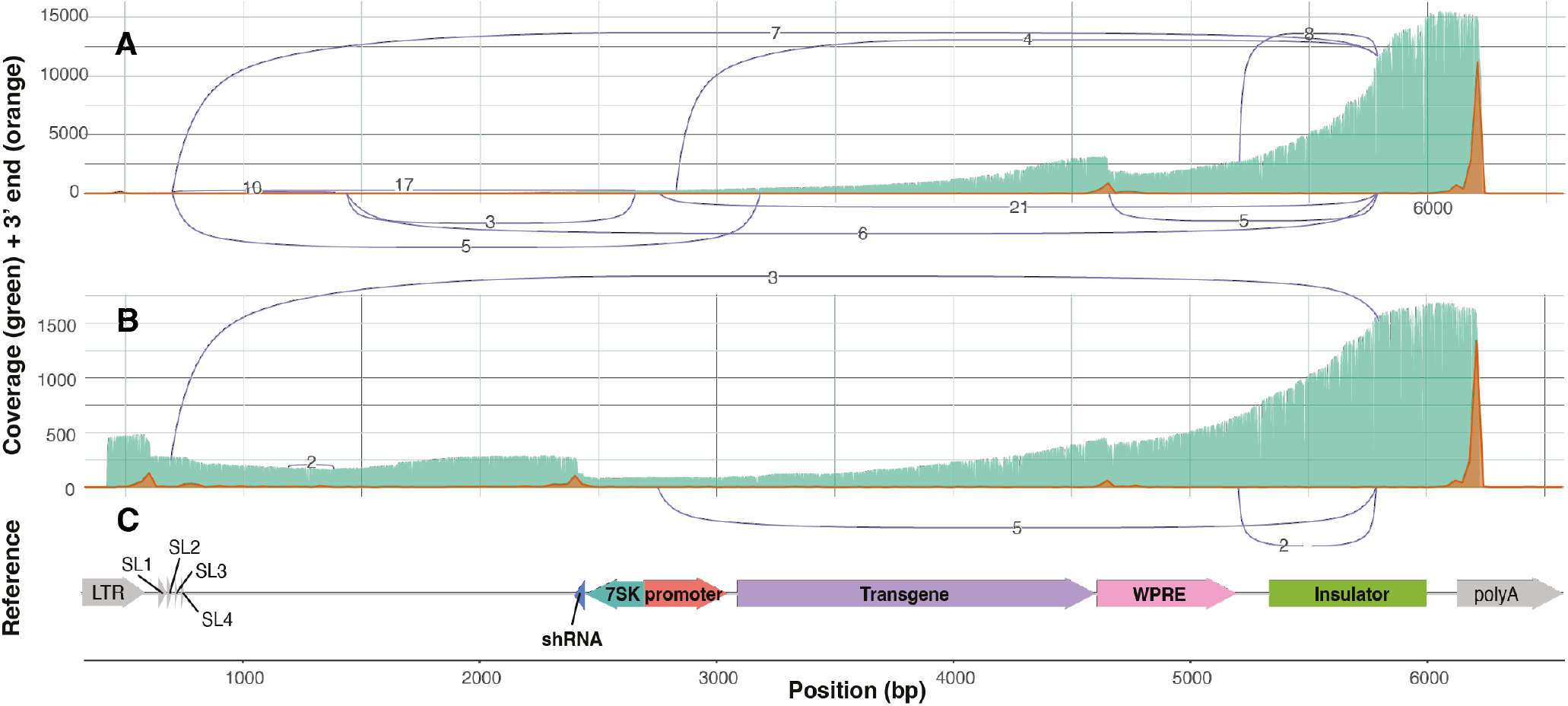
Artificially polyadenylated Nanopore direct RNA sequencing data for a Wiskott-Aldrich Syndrome vector. Plots showing the sequencing coverage (green), location of the 3’ ends of reads (orange) and splicing patterns (purple lines connecting splice donors and acceptors) for Wiskott-Aldrich Syndrome WAS LV1 vectors sequenced by Nanopore direct RNA technology, with and without artificial polyadenylation (performed according to Crabtree et al., 2019 (19)) **(A)** Standard library preparation protocol (no artificial polyadenylation). **(B)** Artificially polyadenylated sequencing data. **(C)** Reference sequence of the Wiskott-Aldrich Syndrome vector, with various features annotated.

### Insights from sequencing can be used to optimise vectors

Having identified the two most significant contributors to incomplete lentiviral RNA in our WAS vectors (the potential cryptic polyA site and hairpin-associated truncations), we sought to improve the vector by modifying these sites. Based on the WAS LV3 vector, a new vector (WAS LV5) was designed, with both the shRNA cassette (a potential Drosha cleavage site responsible for 8.61% of reads being incomplete) removed and the ATTACA motif (a potential cryptic polyA site responsible for approximately 4-10% of reads being incomplete) mutated to ATTTCA. The other potential stem-loop adjacent cleavage site, located in Psi and responsible for 11.26% of reads being incomplete was left unchanged given the important role of this region in nuclear export and packaging of the lentivirus. In addition, a new reverse transcriptase enzyme (Induro) was used to stabilise the RNA during library preparation, due to its advantages when dealing with highly structured and modified RNA. This enabled the generation of longer reads and therefore better coverage of the lentiviral RNA when compared to the previous protocol, further improving the direct RNA sequencing approach (Supplementary Figure 3). Mutation of the cryptic polyA site did not eliminate premature termination but led to a reduction in the percentage of reads terminating in a 150 bp window around this region to 3.39% in WAS LV5, and to 2.04% in WAS LV6 explained below. The mechanism of persistent premature termination is still unclear, and further mutations may be required to prevent it completely. Interestingly, a new and abundant splicing pattern that removes most of the promoter and a small part of the start of the transgene was observed in these modified vectors. 19.35% of WAS LV5 reads are affected by this splicing pattern, which occurs between the promoter and the transgene (Figure 4A). This particular splicing pattern was not observed in any of the previous vector designs (WAS vectors in Figure 2), although the donor site was used with a different acceptor at very low (<1%) levels. Despite this, splice site prediction with MaxEntScan (21) gives the donor and acceptor sites identical scores, irrespective of the vector version (8.15 for the donor site, 10.82 for the acceptor site).). This splicing event could be eliminated in WAS LV6 by the introduction of two point mutations in the promoter (Figure 4B). More specifically, we introduced T-toA point mutations in two “GT” splice donor motifs occurring within a direct repeat region of the promoter. Introduction of only one point mutation in the observed active GT motif would have likely shifted the activity to the splice donor on the other repeat (Figure 4B). Another set of splicing events can be observed towards the 5’ end of vectors WAS LV5 and WAS LV6, likely as a consequence of the improved protocol with increased 5’ end coverage. The majority of those splicing events share a splice donor relevant for the virus life cycle termed the major splice donor (MSD) (22). Previous attempts to mutate this known splice donor resulted in lower RNA expression, possibly due to reduced RNA stability (23). In this study, we did not attempt to mutate any of the virus-related parts of the vector to avoid potential negative consequences. In sum, although the complex nature of lentiviral vectors makes them challenging to optimise, direct RNA sequencing allows for the quality control of packaged lentiviral RNA and facilitates the development of improved vectors.

**Fig. 4.**
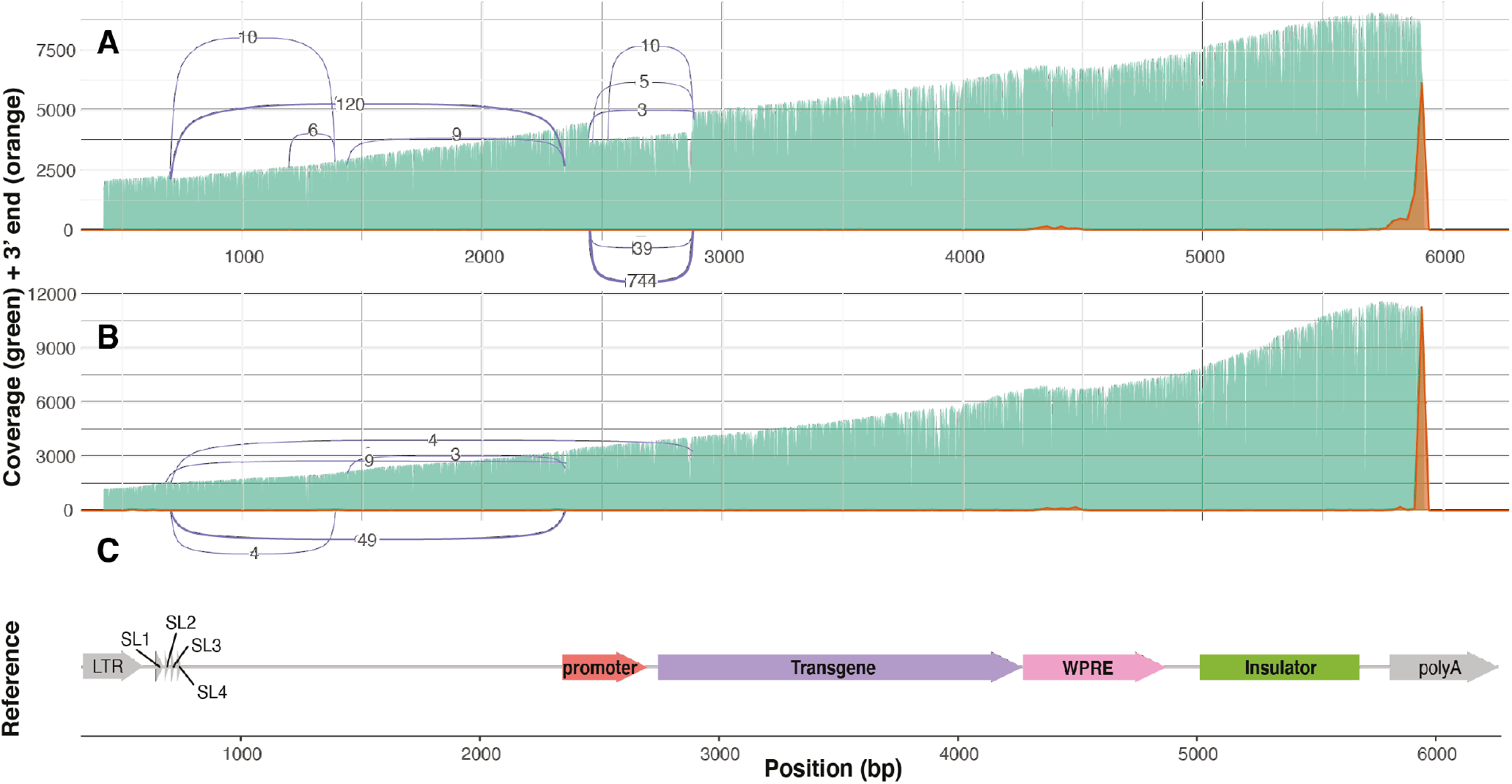
Nanopore direct RNA sequencing of optimised Wiskott-Aldrich Syndrome vectors. Plot showing the sequencing coverage (green), location of the 3’ ends of reads (orange) and splicing patterns (purple lines connecting splice donors and acceptors) for two Wiskott-Aldrich Syndrome vectors sequenced by Nanopore direct RNA technology **(A)** WAS LV5 sequencing data (contains a single A to T mutation in the potential cryptic polyA site). **(B)** WAS LV6 sequencing data (contains an additional two point mutations in the promoter to remove splice donor sites). **(C)** Reference sequence of the Wiskott-Aldrich Syndrome vector, with various features annotated.

## Discussion

Achieving reliable production of intact lentiviral vectors will have a significant impact on their manufacturing and clinical implementation. To this end, we have evaluated the application of three different long-read sequencing technologies in order to determine the most appropriate approach for lentiviral vector QC and to allow improvements to lentiviral vector design. We found Nanopore direct RNA sequencing to be the most suitable, as it, unlike PacBio cDNA sequencing, which generated relatively few reads, provided high sequencing coverage of the vector without any internal priming as observed in the Nanopore direct cDNA data. Using this Nanopore direct RNA sequencing approach, we were able to identify cryptic splice and polyA sites in lentiviral vectors. In order to improve the Nanopore direct RNA sequencing approach, we tried to incorporate an RNase treatment to increase the proportion of vector-aligned reads, artificial polyadenylation to investigate whether there were incomplete RNAs without natural polyA tails and use of the Induro reverse transcriptase to increase read length and thereby coverage. The RNase treatment was unsuccessful, as there was only a slight increase in reads aligning to the vector and RNA degradation rendered the data patchy, due to the decrease in read length (Supplementary Figure 1; Supplementary Table 3). We suspect that this may be due to most of the sequenced human RNA coming from within extracellular vesicles, which are often co-purified with lentivirus and have been shown to contain human RNA from producer cells(24). Applying more rigorous lentiviral purification would therefore likely improve this sequencing assay, by increasing the percentage of on-target reads. Artificial polyadenylation on the other hand produced interesting results, revealing two highly abundant species of incomplete vector RNA, associated with hairpins. We hypothesise that this may be caused by cutting of lentiviral RNA in the producer cell lines by Drosha. Finally, an Induro-based library preparation method (as compared to the standard SuperScript III protocol) delivered significantly longer reads (Supplementary Figure 3). The Induro RT, which has increased processivity and may perform better with the RNA modifications and secondary structure that are present in lentiviral RNAs, is likely better able to support the RNA strand being sequenced and minimise formation of RNA secondary structures that can block the nanopores. This highlights the importance of using reverse transcription to produce an RNA-cDNA hybrid during direct RNA library preparation of lentiviral samples, due to their secondary structure and suggests that future improvements to reverse transcription may benefit this protocol. Using this assay, we identified and mitigated several problematic sites contributing to incomplete lentiviral RNAs. In both WAS LV1 and Globin LV, a splice site in the chromatin insulator sequence was identified, and subsequent vectors with point mutations or inversions were not subject to splicing at this site (Figure 2). We also identified a potential cryptic polyA site with a non-canonical ATTACA motif as a probable cause of truncation in up to 10% reads, which is located in the Woodchuck Hepatitis Virus Posttranscriptional Regulatory Element (WPRE) that is commonly included in lentiviral vectors to increase expression (15). By mutation of the potentially underlying ATTACA polyA signal, we reduced the usage of the cryptic polyA site to about half. Using the artificially polyadenylated sequencing data, we also identified potential Drosha cut sites at two hairpins within the vector sequence which together result in approximately a fifth of all reads being incomplete. Therefore, to improve these vectors, we tried to prevent these truncations by removing the shRNA cassette that contained one of the two hairpins. However, the removal of the shRNA cassette introduced a new splicing pattern nearby that removed the promoter of 20% of reads. Why this splicing pattern only emerges after the removal of the shRNA is unclear, although one possibility is that antiviral activity of Drosha (wherein it binds the stemloop and confers steric hinderance to prevent viral replication (25)) may have been blocking the spliceosome from accessing this site. Although the splicing pattern was subsequently prevented by mutation of the promoter (Figure 4B), this highlights the difficulty of optimising complex lentiviral vector sequences based solely on sequence prediction and design, and the importance of re-assessing their quality by sequencing after modification in order to identify any new problems that may have arisen. The trade-off of using Nanopore direct RNA sequencing as opposed to cDNA sequencing with either Nanopore or Illumina is that the number of reads generated is relatively low, especially when combined with the low percentage of vector-aligned reads. Thus, only around 10,000 lentiviral vector reads are produced on average from a single minION flow cell. Nevertheless, many splice, polyA and hairpin-associated truncation sites can be identified from the sequencing data. Single nucleotide polymorphisms (SNPs), however, are difficult to determine from Nanopore direct RNA sequencing due to the relatively higher rate of both random and systematic errors (26), as well as the abundance of modified bases in lentiviral RNA, which can lead to mis-calling or insertions/deletions during base calling. These issues may be addressed through the recently released Nanopore SQK-RNA004 kit, which promises higher throughput and sequencing accuracy. In summary, Nanopore direct RNA sequencing is a powerful tool for the quality control of lentiviral vectors, allowing for the identification of sites that cause truncations. It can be easily implemented using the affordable, handheld minION device, which allows for rapid, in-house sequencing. To identify all sources of truncation in a vector, including hairpin-associated truncations that are not naturally polyadenylated, we recommend that two Nanopore direct RNA sequencing runs are performed: one with and one without artificial polyadenylation. This will enable rapid analysis and subsequently improvement of lentiviral vectors to increase the proportion of fulllength RNA and hopefully facilitate the manufacturing of better lentiviral vectors in the future.

## Materials and Methods

### Lentiviral vector design

All lentiviral vectors used in this study (listed in Supplementary Table 5) are derived from a pCl20c vector backbone (27) and contain a rabbit β-globin polyadenylation signal (“polyA” in Figures 1-4) downstream of the 3’LTR to provide a stronger polyadenylation signal for the vector transcript. In the Globin LV (Figure 1), gamma globin is expressed from a 254 bp β-globin promoter, placed in reverse orientation with respect to the viral transcript. The expression is regulated by a β-globin locus control region (LCR) region consisting of hypersensitivity sites (HS) 2, 3 and 4 elements. A second expression cassette is inserted in forward orientation and downstream of the globin expression cassette. It expresses an shRNA targeting human HPRT under the control of a human 7SK RNA Pol III promoter. As a safety and anti-silencing element, a 400 bp extended core element of the cHS4 β-globin chromatin insulator is inserted in the 3’LTR in reverse orientation with respect to the viral transcript. All WAS vectors contain a strong synthetic promoter driving the expression of a transgene encoding for human WAS, in forward orientation (relative to the viral transcript). In the 3’ UTR region downstream of the WAS CDS, the vectors contain a Woodchuck Hepatitis Virus (WHV) Posttranscriptional Regulatory Element (WPRE) that harbors 6 point mutations to abrogate WHX truncated protein expression (28). In some WAS vectors, a human 7SK RNA Pol III promoter drives the expression of an shRNA targeting HPRT, in reverse orientation to and located upstream of the first expression cassette. Vector “WAS LV1” contains the original sequence of the 650 bp insulator, reconstructed from the public database (NCBI entry GGU78775) and from the description of the cloning strategy used by Urbinati et al.(16) in reverse orientation. A second vector “WAS LV2” contains two A-to-T point mutations in “AG” splice acceptor motifs in the insulator with the purpose of abrogating splice acceptor activity at these sites. Whereas one mutated “AG” motif corresponds to the known splice acceptor site (12)(13) also present in the Globin LV, the other mutated “AG” motif (specific to the 650 bp and not part of the 400 bp cHS4 insulator variant) was predicted as a likely splice acceptor site in silico (NetGene2(17)). A third vector “WAS LV3” contains three A-to-T point mutations in “AG” motifs in the insulator. In addition to the two “AG” motifs mutated in “WAS LV2”, a third “AG” motif located 10 bp away from the known problematic splice acceptor site (12)(13) is mutated, consistent with another published improved vector with a 400 bp cHS4 insulator variant (13). Vector “WAS LV4” harbors a cHS4 650 bp insulator in forward orientation to de-couple splicing and polyadenylation. Vector “WAS LV5” is derived from WAS LV3 with two introduced changes: the shRNA expression cassette is deleted and the ATTACA motif in the WPRE element is mutated to ATTTCA by introducing a T=>A point mutation.

### Lentiviral production and RNA purification

WAS Lentiviruses were produced by transient transfection of transfer plasmids containing the constructs of interest into GPRTG lentivirus producer cell lines (27)(29). Briefly, 152 ∗106 GPRTG cells were seeded into a 2-STACK culture chamber (Cornig) and transfected 24 h later with 238 µg transfer plasmid using the CalPhos Mammalian Transfection Kit (Takara Bio) according to the manufacturer’s instructions. The transfection mix was replaced with fresh medium 4 h post transfection and virus-containing supernatant was collected 2 days later. After filtration through a 0.22 µM filter, 31 ml of the filtrate was transferred to each of six 40PA ultracentrifugation tubes (Himac), underlayed with 5 ml 20% (w/v) sucrose in 100 mM NaCl, 20 mM HEPES, 1 mM EDTA, and centrifuged for 2 h at 25’000 rpm and 4°C. After centrifugation, the supernatant was discarded and the viral pellet was resuspended in 50 µl X-Vivo10 medium (Lonza) supplemented with 2% human serum albumin. Resuspended viral pellets were filtered through a 0.22 µM filter, pooled and stored at 70°C. For subsequent analysis by sequencing, lentiviral RNA was purified using the QIAamp Viral RNA Mini Kit (Qiagen) according to the manufacturer’s instructions. Quality control of extracted RNA was performed using a fragment analyzer.

### Artificial polyadenylation

Artificial polyadenylation was carried out based on Crabtree et al. (19) Reactions containing 15 µL of RNA (500 ng), 2 µL of polyA buffer, 2 µL of 10 mM ATP (NEB P0756S), 1 µL of E. coli Poly(A) Polymerase (NEB M0276S) and 0.5 µL of Murine RNase Inhibitor (NEB M0314S) were incubated for 30 minutes at 37°C, 20 minutes at 65°C and then 5 minutes at 98°C before being placed on ice. The RNA was then cleaned up with magnetic beads at 1.8X ratio and eluted in 15 µL nuclease free water ready for the Nanopore direct RNA sequencing protocol. Given the poorer quality of artificially polyadenylated reads, it might be beneficial to adopt a more gentle approach (30).

### PacBio cDNA sequencing

PacBio SMRTbell libraries were prepared according to the manufacturer’s instructions and sequenced on the PacBio Sequel platform with v3.0 chemistry. The generated subreads were demultiplexed and CCS reads were obtained using v4.2.0 of PacBio’s CCS algorithm with default parameters.

### Nanopore direct cDNA sequencing

Nanopore direct cDNA sequencing libraries were prepared using the SQKDCS109 kit, according to the manufacturer’s instructions. Sequencing was performed using a minION R9.4.1 flow cell (FLO-MIN106D) on a gridION instrument.

### Nanopore direct RNA sequencing

Nanopore direct RNA sequencing libraries were prepared using the SQK-RNA002 kit, according to the manufacturer’s instructions. For the final set of experiments (Figure 4), the SuperScript III RT used in the reverse transcription step of protocol was replaced with Induro RT. Reactions containing 2 µL of dNTP, 8 µL of 5X Induro RT Buffer (NEB M0681S), 12.6 µL nuclease free water, 0.4 µL RNase Inhibitor and 2 µL Induro RT (200 U/µL; NEB M0681S) were incubated for 15 minutes at 60°C, 10 min at 70°C and then cooled to 4°C immediately. Sequencing was performed using minION R9.4.1 flow cells (FLO-MIN106D) on a gridION instrument.

### Bioinformatic analysis

Following basecalling and basic sequencing quality control using the seqkit (31) stats function, reads were aligned to both the lentiviral vector and human genome reference sequences using minimap2 (32) in splice mode. The samtools (33) software was used to determine alignment statistics (samtools flagstat) and coverage of the vector reference sequences (samtools depth). Positions of 3’ read endings were extracted from the PAF output file generated by minimap2 (32) in R (34) and sashimi plots of the splicing patterns were generated using ggsashimi (35). This information was then collated in R (34) and plotted with ggplot2(36), using the gggenes (37) package to plot the vector reference sequence, ggrepel (38) to label the vector elements and cowplot (39) to combine multiple plots into a single panel. Cryptic polyA sites in the data were predicted based on the motifs described in the polyAsite 2.0 database (40), and hairpins were modelled using the RNAfold (41) webserver. A detailed bioinformatic tutorial can be found at https://kzeglinski.github.io/lentiviral_qc/bioinf.html

## ACKNOWLEDGEMENTS

We thank Esther Chen for virus production of the Globin LV, Martina Biserni for support with lentivirus production and RNA extraction, Alban Ramette and team at IFIK for the Nanopore sequencing service, Genewiz For the PacBio sequencing service and the CSL/WEHI Translational Data Science Alliance for funding this project.

## Declaration of interests

One or more of the model vectors used as the basis for this study were published in International Publication Nos. WO2019018383, WO2020139796 or WO2022232191. Authors C.M., C.V. and F.A. are named as inventors. K.Z and Q.G have previously received travel and accommodation expenses from Oxford Nanopore Technologies.

## Data availability statement

Sequencing data for this study is available from the European Nucleotide Archive (PRJEB72210)

## Author contributions

KZ: Conceptualisation, Data Curation, Formal Analysis, Visualisation, Writing-Original draft preparation, CM: Conceptualisation, Investigation, Writing-Review & Editing, MER: Formal Analysis, Supervision, MA: Supervision, CV: Supervision, Resources, RB: Supervision, MJ: Investigation, QG: Conceptualization, Formal Analysis, Supervision, FA: Conceptualisation, Resources, Writing- Review & Editing, Project administration, AH: Conceptualisation, Writing-Review & Editing, Supervision

## Notes

https://www.ebi.ac.uk/ena/browser/view/PRJEB72210

https://kzeglinski.github.io/lentiviral_qc/bioinf.html

### Summary of Updates

Updated Figure 4 and accompanying text to include new data; added links to analysis code and raw sequencing data

