## Supplementary for "An optimised protocol for quality control of gene therapy vectors using Nanopore direct RNA sequencing"

### Supplementary Note 1: RNase treatments do not increase the proportion of vector-aligned reads

As the percentage of vector aligned reads is relatively low (particularly in the WAS samples), we investigated the use of an RNase treatment to degrade human RNAs while the lentiviral RNA remained protected inside the virus. This slightly increased the percentage of reads aligning to the vector reference sequence (3.21% vs 2.01% in the control; Supplementary table 3), although because the median read length of RNase treated reads (425 bp) was much shorter than the control (722 bp) there was little difference in the number of bases aligned to the vector.

Sequencing coverage of the RNase treated sample (Supplementary figure 1B) displayed a much stronger 3' to 5' decay when compared to the control (Supplementary figure 1A) which agrees with the shorter median read length (Supplementary table 3). As a result, although the overall pattern of peaks (both 3' end and sequencing coverage peaks) was the same, due to low coverage at the 5' end, the RNase treated sample fails to capture many of the splicing events that occur in that region.

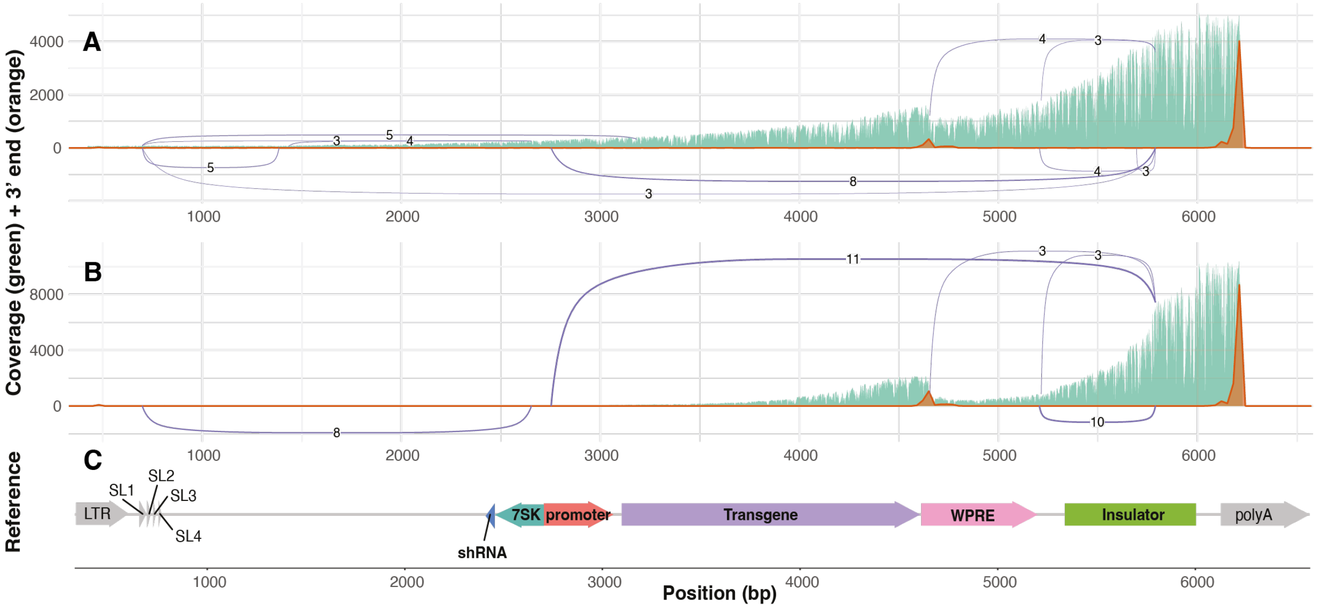


**Fig. 1.** Sequencing coverage (green), location of the 3' ends of reads (orange) and splicing patterns (purple lines connecting splice donors and acceptors) for Wiskott-Aldrich Syndrome vectors sequenced by Nanopore direct RNA sequencing, with and without RNase treatment **(A)** Control sequencing data (no RNase treatment). **(B)** RNase treated sequencing data. **(C)** Reference sequence of the Wiskott-Aldrich Syndrome vector, with various features annotated.

### Supplementary Note 2: Artificial polyadenylation produces consistent results for both WAS LV1 and 2

Similar to the results for WAS LV1, the artificially polyadenylated sample from WAS LV2 shows a similar pattern to the normal (non-polyadenylated) data (green, Supplementary figure 2). 4.19% of all reads terminate at the potential cryptic polyA site in the WPRE, while 11.66% terminate around the location of the shRNA and 5.53% in the 5' LTR of the vector. This is in contrast to WAS LV1, where more reads terminated in the 5' LTR (11.26%) than at the shRNA site (8.61%) which may suggest that the mechanisms behind lentiviral vector truncation have batch-to-batch variability. The estimated percentage of full-length RNA was similar to that of WAS LV1 (60-78% and 60-75%, respectively), and no splicing was detected.

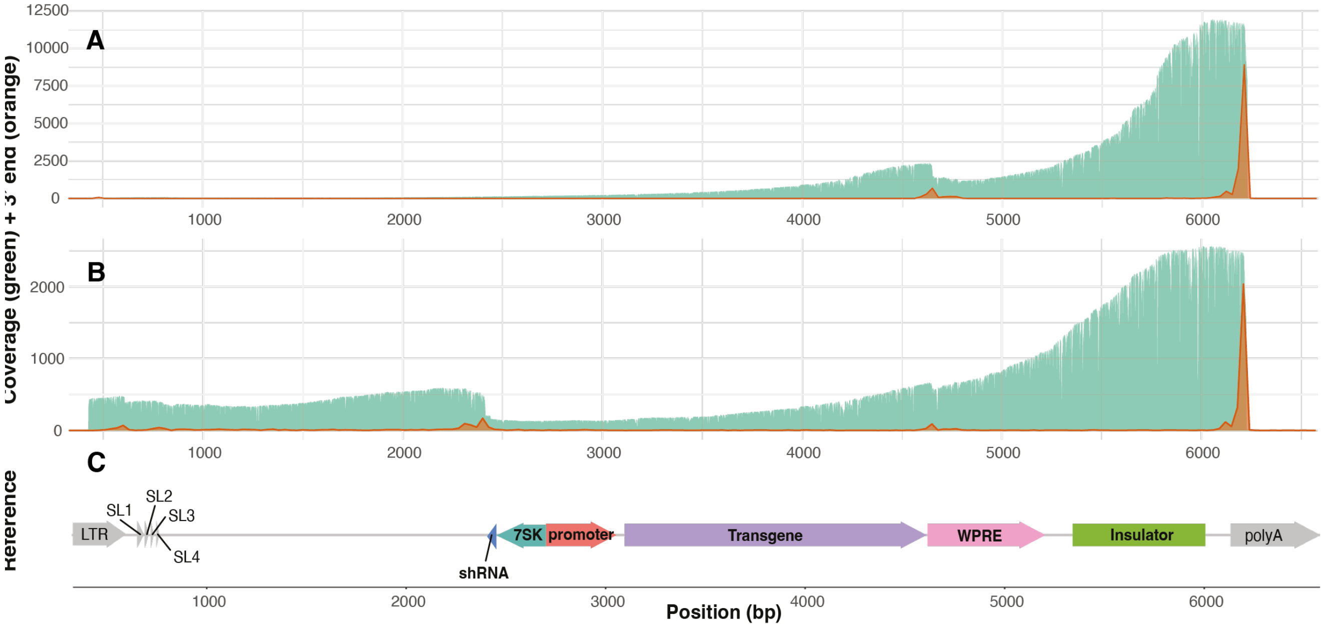


**Fig. 2.** Plot showing the sequencing coverage (green) and location of the 3' ends of reads (orange) for Wiskott-Aldrich Syndrome WAS LV2 vectors sequenced by Nanopore direct RNA technology, with and without artificial polyadenylation **(A)** Standard library preparation protocol (no artificial polyadenylation). **(B)** Artificially polyadenylated sequencing data. **(C)** Reference sequence of the Wiskott-Aldrich Syndrome vector, with various features annotated.

### Supplementary Note 3: Induro reverse transcriptase optimises the sequencing protocol by generating longer reads

In order to improve the read length from the Nanopore direct RNA sequencing approach and thus reduce the 3' coverage bias observed (Supplementary figure 3A), we trialled a new library preparation method using the Induro reverse transcriptase. Although only the RNA is sequenced, synthesising a strand of cDNA is an important step in Nanopore direct RNA sequencing library preparation as it helps to increase throughput, likely through the prevention of RNA secondary structure formation (which may block translocation through the Nanopore). The Induro reverse transcriptase has been shown to generate longer reads, improving the throughput and 5' coverage in Nanopore direct RNA sequencing. Thus, we compared the conventional Nanopore direct RNA sequencing approach (using SuperScript III) to the improved method (using Induro) to determine whether such increases would be beneficial for the quality control of lentiviral RNA. Despite coming from the same input RNA, reads in the Induro sample (Supplementary figure 3B) were much longer than those from the conventional approach (Supplementary figure 3A), with coverage at the 5' end being much higher. This reveals additional, although rare, splicing patterns at the 5' end of the vector, as well as making it easier to visually identify more abundant splicing patterns (Supplementary figure 3B). However, the percentage of reads involved in the most common splicing pattern (calculated as the number of spliced reads divided by the total number of reads at that position) is similar for both reverse transcriptases (23.91% for SuperScript III vs 19.35% for Induro).

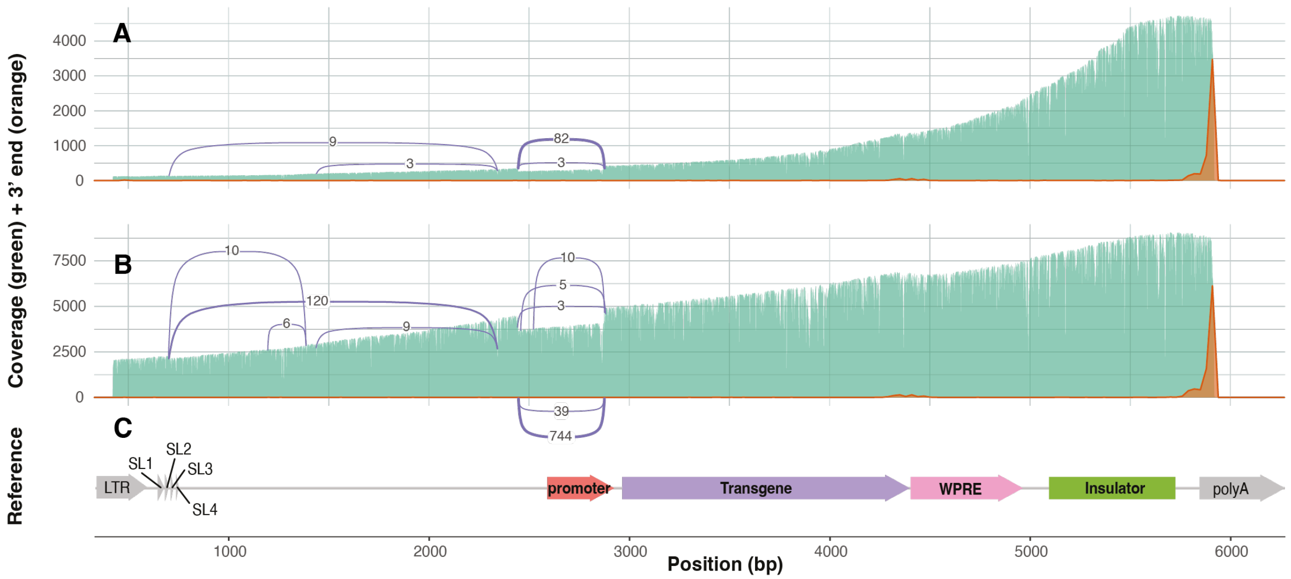


**Fig. 3.** Sequencing coverage (green), location of the 3' ends of reads (orange) and splicing patterns (purple lines connecting splice donors and acceptors) for the same RNA from Wiskott-Aldrich Syndrome lentiviral vectors sequenced by Nanopore direct RNA sequencing with different reverse transcriptases **(A)** Control sequencing data (using the standard protocol with SuperScript III reverse transcriptase). **(B)** Improved protocol sequencing data (using a modified protocol with Induro reverse transcriptase) **(C)** Reference sequence of the Wiskott-Aldrich Syndrome vector, with various features annotated.

### Supplementary Note 4: Tables

**Table 1.** Alignment statistics for long-read RNA sequencing technology comparison

| Sequencing technology | Number of reads | Percentage of reads mapped to vector | Percentage of reads mapped to human genome | Median read length (bp) |
|---|---|---|---|---|
| Nanopore direct RNA | 273,029 | 7.97% | 90.71% | 701 |
| Nanopore direct cDNA | 491,270 | 17.18% | 82.40% | 980 |
| PacBio cDNA | 50,159 | 6.71% | 83.37% | 1,795 |

**Table 2.** Alignment statistics for Nanopore direct RNA sequencing data of WAS vectors

| Sample | Number of reads | Percentage of reads mapped to vector | Percentage of reads mapped to human genome | Median read length (bp) |
|---|---|---|---|---|
| WAS LV1 | 743,182 | 2.44% | 96.18% | 630 |
| WAS LV2 | 445,076 | 3.12% | 96.35% | 709 |
| WAS LV4 | 621,958 | 2.04% | 96.99% | 644 |
| WAS LV3 | 489,502 | 1.47% | 97.99% | 712 |
| WAS LV5 | 205,061 | 4.95% | 94.19% | 918 |
| WAS LV6 | 314,174 | 4.25% | 85.95% | 761 |

**Table 3.** Alignment statistics for Nanopore direct RNA sequencing data of RNase treated sample and control

| Sample | Number of reads | Percentage of reads mapped to vector | Percentage of reads mapped to human genome | Median read length (bp) |
|---|---|---|---|---|
| RNase treated | 407,517 | 3.21% | 96.05% | 425 |
| Control | 304,079 | 2.01% | 98.00% | 722 |

**Table 4.** Alignment statistics for Nanopore direct RNA sequencing data of artificially polyadenylated samples

| Sample | Number of reads | Percentage of reads mapped to vector | Percentage of reads mapped to human genome | Median read length (bp) |
|---|---|---|---|---|
| PolyA treated WAS LV1 | 476,115 | 0.61% | 96.33% | 295 |
| PolyA treated WAS LV2 | 380,551 | 1.13% | 96.36% | 335 |

**Table 5.** Summary of vectors used in this study.

| LV name | Transgene | shRNA | Insulator | WPRE and promoter |
|---|---|---|---|---|
| Globin LV | Gamma globin | yes | cHS4 400bp | WPRE with 6 point mutations |
| WAS LV1 | WAS | yes | cHS4 650bp | WPRE with 6 point mutations |
| WAS LV2 | WAS | yes | cHS4 650bp with two point mutations | WPRE with 6 point mutations |
| WAS LV3 | WAS | yes | cHS4 650bp with three point mutations | WPRE with 6 point mutations |
| WAS LV4 | WAS | yes | cHS4 650bp inverted | WPRE with 6 point mutations |
| WAS LV5 | WAS | no | cHS4 650bp with three point mutations | WPRE with 6 point mutations plus another point mutation to change the ATTACA motif to ATTTCA |
| WAS LV6 | WAS | no | cHS4 650bp with three point mutations | WPRE with 6 point mutations plus another point mutation to change the ATTACA motif to ATTTCA and two point mutations in the promoter to remove splice donor sites |
